# The fitness landscape of the mobilized colistin resistance gene *mcr-1*

**DOI:** 10.1101/2022.07.21.500813

**Authors:** Ziyan Guo, Siyuan Feng, Lujie Liang, Jiachen Li, Lan-Lan Zhong, Hui Zhao, Guo-bao Tian, Jian-Rong Yang

## Abstract

The emergence and global discovery of the mobilized colistin resistance gene, *mcr-1*, limits the clinical effectiveness of colistin as last-resort antibiotic against multiple-drug resistant bacteria. Although MCR-1 exhibits low levels of colistin resistance in *E. coli*, it is currently unknown whether *mcr-1* may potentially evolve a high level of colistin resistance in the future. Here, we generated *mcr-1* mutant library that included 27,965 variants, covering 94.44% of the single-nucleotide mutations. We quantified the relative growth of these variant strains under colistin exposure. Notably, most mutations in the catalytic domain, but not in the transmembrane domain, reduce colistin resistance. The structural basis for the mutational effects revealed that the transmembrane domain may contribute to MCR-1 evolution. Altogether, our data shed light on the prediction of MCR-1 evolution, which could be useful for designing novel treatment strategies.

## Main

The emergence and global discovery of the first mobilized colistin resistance gene, *mcr-1*, threatens the clinical effectiveness of colistin as last-resort antibiotic against multiple-drug resistant bacteria, especially carbapenem-resistant Enterobacteriaceae^1,2^. It is generally accepted that heavy use of colistin as a growth promoter in veterinary applications correlates with the emergence and spread of *mcr-1*^3,4^. Several recent studies have demonstrated a reduction in prevalence of *mcr-1* since May 1, 2017 when China banned the use of colistin as an additive in animal feed^5-7^. However, we found, using a time-series analysis of inpatient colonization data from 2011 to 2019, that there is still a low level of *mcr-1* prevalence among inpatients^8^, which may be related to the approval of colistin for human clinical use in China in January 2017^8^. Notably, previous research demonstrated that the co-existence of *mcr-1* and carbapenemase genes increased after introduction of polymyxin into clinical practice^9^. Therefore, the risk of transmission of *mcr-1* in clinical settings, particularly that of infection by multidrug-resistant *mcr-1* positive bacteria, should not be underestimated, despite the decreased prevalence.

The *mcr-1* gene is a member of the phosphatidylethanolamine (pEtN) transferase family and transfers a pEtN residue to the lipid A present in the outer membrane of Gram-negative bacteria^10^. Such modified lipid A neutralized the negative charge and decreased binding of the positively charged colistin, allowing colistin resistance. Several reports have shown the relevance of MCR-1 expression on bacteria fitness^11-13^ which was caused by the embedding of MCR-1 in outer membrane and the modification of lipid A^14^. More recently, we further demonstrated that MCR-1 causes lipid remodeling that resulted in an outer membrane permeability defect, thus compromising the viability of Gram-negative bacteria^15^.

Although ten mobilized colistin resistance genes referred to as *mcr-1* to *mcr-10* had been identified^16^, *mcr-1* is the most predominant type and conferred relatively high level of colistin resistance among MCR family members (MIC=4 or 8μg/ml)^17^. The potential evolution towards even higher level of colistin resistance (HLCR) in mcr-1-positve strains is of great concern. Several different groups have investigated how *mcr-1* might impact emergence of HLCR mutants^18,19^. Using *in vitro* stepwise colistin exposure, three groups reported that MCR-1 expression were anticorrelated with the development of HLCR in *E. coli* and *Klebsiella pneumoniae*. However, another study reported that MCR-1 conferred increased mutation rate in *E. coli* but did not affect the HLCR mutation rates in *K. pneumoniae*^20^. There is an ongoing debate about the effects of MCR-1 on the development of HLCR in Gram-negative bacteria. Although 31 *mcr-1* variants had been identified from clinical isolates^21^, it is currently unknown whether *mcr-1* may potentially evolve high level of colistin resistance.

To answer this question, we generated all possible single mutation of *mcr-1* and measure their fitness effects in competitive experiment, thus assessing the local fitness landscape^22,23^ of *mcr-1*. Adapting our previous method^24^, we generated a comprehensive *mcr-1* variant library, in which each variant contained a unique 30 nt barcode in the downstream region of the *rrnB* terminator (Figure 1A). These variants were cloned into a low-copy-number plasmid and ∼400, 000 colonies were generated (**See Methods**). To determine the correspondence between the barcodes and genotypes, we used the PacBio Sequel System to determine both the 1626-bp *mcr-1* sequence and the 161bp barcode + terminator sequence for a single molecule. Finally, we captured at least one barcode for a total of 171,769 variants, covering 99.96% of the single-nucleotide mutations (Figure S1A). Next, competitive experiments were performed under colistin selection (0.125, or 0.25 × minimal inhibitory concentration, or MIC) and the variant frequency was estimated via NovaSeq sequencing of barcodes. Genotype frequencies were highly correlated between two T0 technical repeats (Pearson’ s correlation, R ≥ 0.9999) (Figure S1B). To accurately estimate the relative growth, ∼27900 genotypes with read counts ≥ 100 at T0 were included for further analyzed. Among these genotypes, 4590/4860 (94.4%) possible single mutations were represented (Figure 1B). Genotype frequencies were highly correlated between biological replicates at T1 (Pearson’ s correlation, R ≥ 0.9999, OD_600_=0.7) (Figure 1C).

**Fig. 1.**
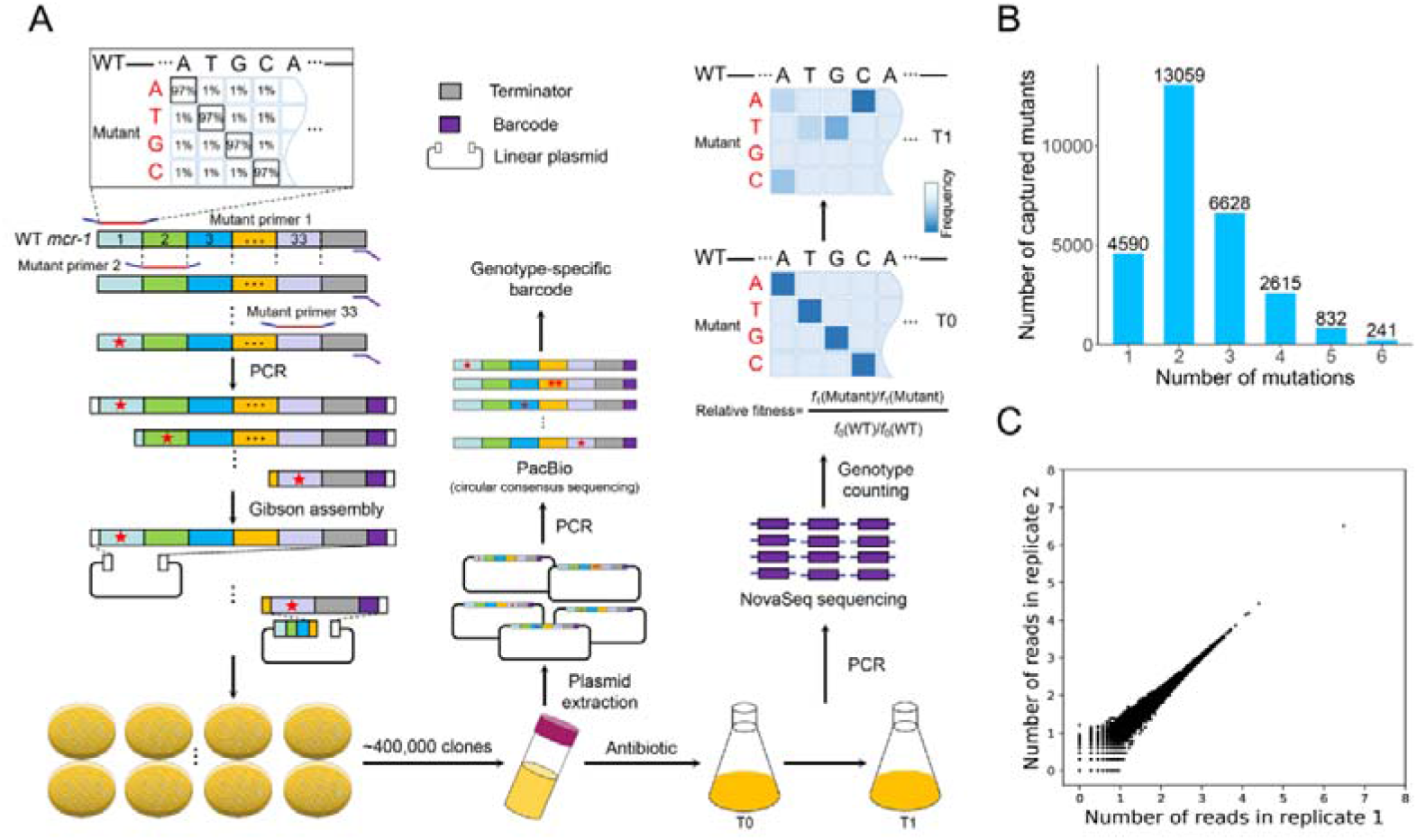
Determining the fitness landscape of *mcr-1* gene. **(A)** Illustration of the experimental workflow for assessing the fitness landscape of MCR-1. The *mcr-1* variant library was generated using mutant primers (3% per-site mutation rate). Then, the variant library was cloned into the low-copy-number plasmid by Gibson assembly. The plasmid library was sequenced with the PacBio Sequel after plasmid extraction in order to determine the correspondence between genotypes and barcodes. Competition experiments were conducted in LB liquid medium containing colistin. When required, the bacterial culture was collected and the plasmid library was extracted for library construction. After barcode amplification, Illumina NovaSeq sequencing was used to obtain the frequency of mutant genotype *f* (Mutant) or wild-type *f* (WT). The relative growth of each genotype was evaluated as the increase in frequency under antibiotic selection relative to wild type *mcr-1*. **(B)** Numbers of variants with 1, 2, 3, 4, 5, and 6 single-nucleotide mutations whose relative growth was determined in our experimental pipeline. **(C)** Comparison of genotype frequencies between biological replicates at T1 in the presence of 0.25 x MIC colistin.

Upon closer examination of the fitness landscape of *mcr-1*, we found that the most beneficial mutation in 0.25 × MIC colistin grows 3.78-fold faster than wildtype *mcr-1* (Table S1-3. Figure 2 A). Such an effect is much smaller than previous observations made in another antimicrobial resistance gene *bla*_*CTX*-M-14_ (encoding beta-lactamase), whose mutation exhibits over 4,000-fold increase in relative growth upon ceftazidime exposure compared to that of wildtype^24^. In addition, the result of antimicrobial susceptibility testing showed that there was no significant difference between wildtype MCR-1 and mutations with increased relative growth (Table S4, Figure 2B). Additionally, most mutations in the catalytic domain (nucleotide positions 600-1626), but not in the transmembrane domain (nucleotide positions 1-600), were deleterious. Indeed, when we overlaid the maximum relative growth observed for the substitutions of each amino acid on the three-dimensional structure of the wild-type MCR-1 (Figure 2C and 2D), we again noticed more beneficial mutations in the transmembrane domain. Interestingly, our recent work demonstrated that a potential lipid A binding pocket of MCR-1 was essential for colistin resistance and maintaining bacterial viability^15^. These observations suggest that MCR-1 variants, at least all possible single nucleotide substitutions, could not lead to a substantial increase in colistin resistance. To our knowledge, this is the first comprehensive evaluation of the phenotypic effects of almost all possible single-nucleotide mutations within the coding sequence of the *mcr-1* gene, which provides the strongest evidence for the limited evolvability of *mcr-1*.

**Fig. 2.**
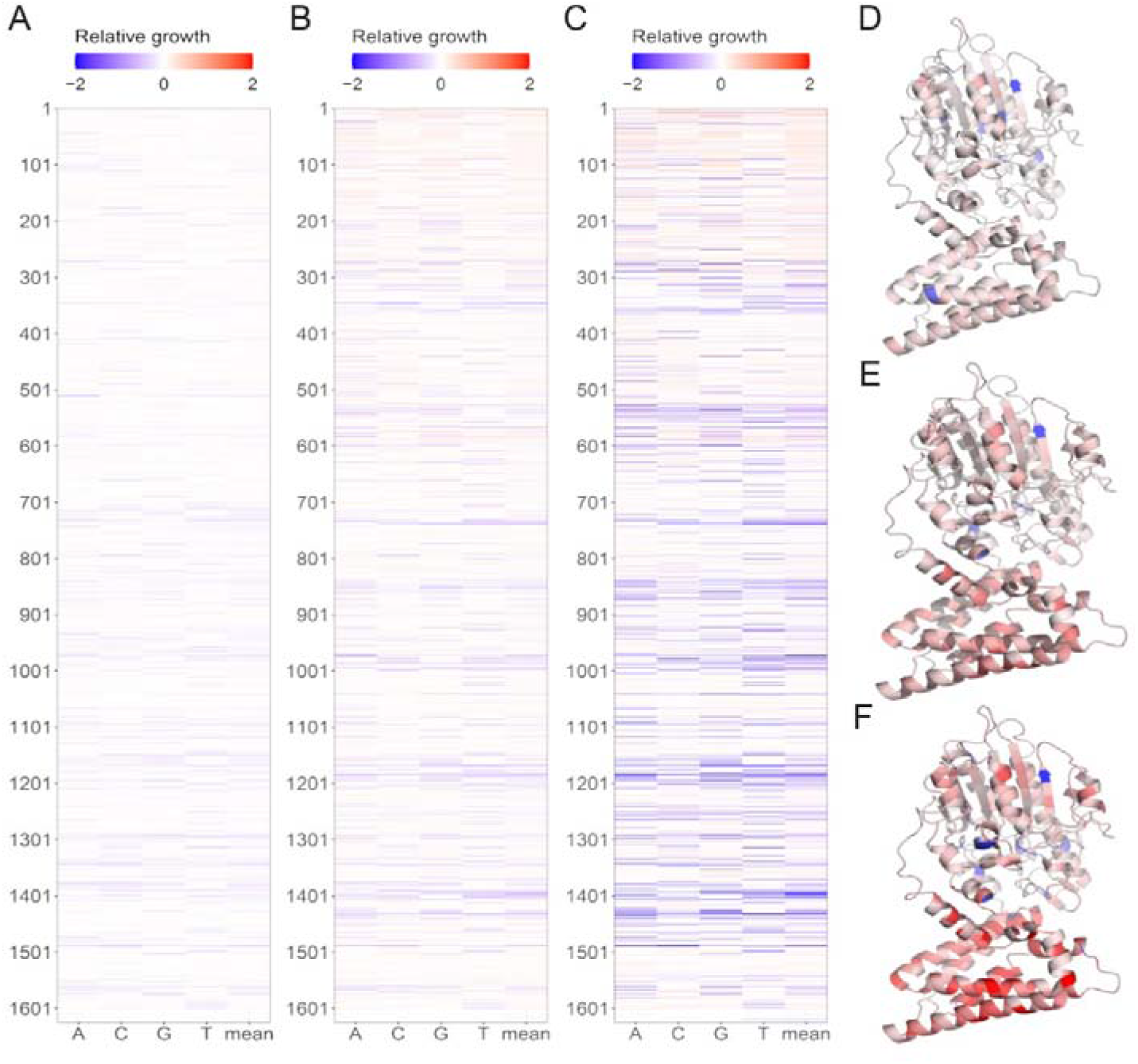
Fitness effects of all single-nucleotide mutations in MCR-1 under colistin selection. (**A**-**B**) Fitness landscape of *mcr-1* in the presence of 1/8 x MIC **(A)** or 1/4 x MIC **(B)** of colistin at T1. Each tile represents a variant with one single-nucleotide mutation (*x* axis) at one specific position (*y* axis), whose relative growth is indicated by the color of the tile scaled according to the corresponding color scale bar on top. (**C**-**D**) For the relative growth measured in the presence 1/8 x MIC (**C**) or 1/4 x MIC (**D**) of colistin, the maximum value of relative growth associated with the nine single-nucleotide substitutions at an amino acid position is indicated as colors overlaid on the three-dimensional structure of MCR-1.

## Materials and methods

### Construction of the plasmid library of *mcr-1* variants

To obtain a template for amplification of *mcr-1* variant and plasmid backbone, we generated a backbone containing the expression cassette of *mcr-1* and pACYCDuet-1, a low-copy chloramphenicol-resistant plasmid. Briefly, the template plasmid was constructed via two rounds of PCR amplification. Phanta Max Super-Fidelity DNA Polymerase (Vazyme, China) was used in all amplification reactions. First, the *mcr-1* gene with the arabinose-regulated ParaB promoter was amplified from pBAD24-mcr-1^25^, and the *rrnB* terminator was amplified from the integrated pOSIP-KH plasmid. After the purification of the PCR product, the two DNA fragments were concatenated by fusion PCR to obtain the full expression cassette of *mcr-1*. pACYCDuet-1 was digested with restriction enzyme FastDigest *Xba*I and FastDigest *Xho*I (Thermo Scientific, America) according to manufacturer’ protocol. Restricted DNA products were analyzed by gel electrophoresis, and the target bands were purified using QIA quick PCR purification kit (Qiagen). The expression cassette was subsequently cloned into digested pACYCDuet-1. Positive colonies were confirmed by PCR and Sanger sequencing. The resulting plasmid was named pACY-Para-mcr-1 and stored at −20°C. All the primer used in this study was listed in Table S5.

Doped oligonucleotides were synthesized by IDT (https://www.idtdna.com/) as previously described^24^. Since the length of chemically synthesized oligonucleotides with degenerate sites containing manually defined nucleotide fractions was limited to 90 nt and the invariant regions at both ends of the oligonucleotides were required for PCR, we designed only 50 variable sites for each oligonucleotide, and the leading and trailing 20 nt were invariant sequences identical to those of wild-type *mcr-1*. In addition, for the doped oligonucleotides, each position contained the wild-type nucleotide at a 97% frequency and a 1%:1%:1% mix of the other three nucleotides. As a result, the library exhibited a mutation rate of approximately 3% per nucleotide. According to theoretical and previous empirical data^26^, it was expected that ∼25% of the synthesized oligonucleotides were wild type, while 25% contained single-nucleotide mutations and the others contained multiple mutations. To acquire variant library for *mcr-1* with 100% coverage for all possible single-nucleotide mutations, a total of 33 mutant primers was synthesized to cover the full length of *mcr-1* (1626bp).

The gene variant library was constructed via one-step cloning following the manufacturers’ instructions of the commercial kits unless otherwise specified. Briefly, 33 mutant fragments were individually amplified using each mutant primer and Ter-15BC-template-15BC-PstI-R by standard PCR. Notably, Ter-15BC-template-15BC-PstI-R primer containing a 20bp region identical to the 5’ end of *rrnB* terminator and a 30bp region of fully degenerated (randomized) nucleotides serving as a barcode and a 20-bp region identical to linear pACY-Para-mcr-1. Thus, there are 4^30^ ≈ 10^18^ possible sequencing combinations for a 30bp barcode, which is 1000× the number of template DNA molecules used for PCR. After 30 cycles, the PCR products were analyzed by gel electrophoresis, and the target bands were purified using QIA quick PCR purification kit (Qiagen). Next, 33 different plasmid backbones of different lengths were individually amplified from pACY-Para-mcr-1. After 20 cycles, the PCR products were treated with FastDigest *Dpn*I and purified using Cycle Pure Kit (OMEGA, America). Ligation was performed a 3-to-1 ratio in terms of mutant fragment versus its corresponding plasmid backbone, using a Seamless Cloning Kit (Beyotime, China). The product was transformed into BW25113 competent cells and selected on chloramphenicol (30μg/ml). Each mutant fragment contains 50bp mutated region, which leaves 50×3=150 total possible single point mutation. We aimed to collect ∼12,000 clones for each mutant fragment, which is 20× the number of single mutation (150/25% contained single-nucleotide mutations) ×20=12000). In total, the *mcr-1* library comprised approximately 4×10^6^ clones. All colonies were collected from LB agar plates by washing with LB liquid medium. Pooled plasmid variants were extracted with a Plasmid Midi Kit (OMEGA, America) and stored at −80°C.

### Site-directed mutagenesis

To assess the accuracy of the relative growths measured, 5 variants of *mcr-1* were also constructed by one-step cloning. Briefly, two adjacent mutated target fragments with 20-15 bp homologous sequences were amplified by PCR. The PCR products were gel purified using a gel extraction kit (OMEGA). The pACYCDuet-1 plasmid was digested with FastDigest *Xba*I and FastDigest *Xho*I at 37°C for 1 h. Ligation was performed at a vector: insert ratio of 1:3 using a Seamless Cloning Kit (Beyotime, China). The product was transformed into *E. coli* BW25113 competent cells and selected on chloramphenicol (30μg/ml). Recombinant plasmids were purified, and the corresponding *mcr-1* variant was sequenced to confirm the mutation.

### Construction of the *E. coli* BW25113 strain pool

To construct the *mcr-1* variant strain pool, the plasmid library was transformed into *E. coli* BW25113 chemically competent cells. Immediately after heat shock, 1 ml SOC medium was added, and the culture was recovered at 37°C for 1 h. Subsequently, LB agar plates containing 25 mg/ml kanamycin were incubated at 37°C for 12 h. Over 400,000 colonies were collected from LB agar plates by washing with LB liquid medium. *E. coli* BW25113 chemically competent cells were prepared through a modified Hanahan method^27^. Briefly, a single colony from a fresh plate of the strain was inoculated into 2 ml LB medium and cultivated at 37°C at 300 rpm overnight as a seed culture. One milliliter of the seed culture was transformed into 100 ml of LB liquid medium and cultivated at 37°C until the OD_600_ reached a value of 0.4-0.6. Each 25 ml culture was transferred to a chilled 50 ml centrifuge tube and incubated on ice for 15 min. The cell pellet was spun down at 4°C (4000 rpm for 10 min), and the supernatant was discarded. The cell pellet was then resuspended in 30 ml of ice-cold 0.1 M CaCl_2_-MgCl_2_ 280 mmol/l MgCl_2_ and 20 mmol/l CaCl_2_ solution, followed by incubation on ice for 30 mins. After being spun down again 4000 rpm at 4°C for 5 min, the cell pellet was resuspended in 2 ml of iced 0.1 M CaCl_2_-15% glycerol, and 100 μl aliquots of the suspension were transferred to 1.5 ml microtubes and stored at −80°C.

### Competition experiments

After harvesting the *E. coli* BW25113 strain pool, 100 μl aliquots (∼2×10^8^ CFU) of the strain pool were added to 100 ml of LB liquid medium containing 2 μg/ml (0.125 × MIC), 4 μg/ml (0.25 × MIC) colistin. Three replicate competition experiments were performed. To dynamically examine the change in the phenotype, 20 ml of each sample was collected at OD_600_ =0.7.

### Library preparation

For PacBio sequencing, the plasmid library was used as the template, and 25 cycles of PCR were performed to amplify the expression cassette, including the barcode. The PCR product was run on an agarose gel and purified with a gel extraction kit (OMEGA).

For NovaSeq sequencing, plasmid DNA was extracted from the sample of interest. Two rounds of PCR were performed to amplify the barcode from the plasmid library. In brief, 20 cycles of PCR were performed to amplify the barcode-containing fragment, and the purified product was used as the template for the second round of PCR. Twenty-five additional cycles of PCR were then performed to amplify the barcode sequence using primers that also added Illumina TruSeq adapters.

### Antimicrobial susceptibility testing

The MICs of cefotaxime and ceftazidime for *E. coli* BW25113 carrying the *mcr-1* or *mcr-1* mutants were determined using the agar dilution method and interpreted using breakpoints defined by the Clinical and Laboratory Standards Institute (CLSI).

### Associating barcodes and *mcr-1* genotypes via PacBio sequencing

We used three single-molecule real-time (SMRT) cells on the PacBio Sequel platform to sequence the constructed plasmid pool and obtained a total of 3.5×10^7^ raw subreads (Fig. S1A). There was a nonnegligible probability of the presence of heterologous dsDNA molecules since the ssDNA molecules of different mutants in the plasmid pool were highly similar and therefore capable of forming heterologous duplexes. To avoid base calling errors caused by heterologous dsDNA, we used BLASR with default parameters^28^ to map all subreads of each zero-mode waveguide (ZMW) to the wild-type sequence of *mcr-1* and divided them into positive and negative strands. We then used the CCS algorithm (*--min-length 900 --max-length 1600 --min-passes 5*)^29^ to call consensus sequences separately from the subreads derived from positive and negative strands, with at least five subreads each, which corresponded to a base-calling error rate ≤1% (Fig. S1B). From each CCS without any indels, we extracted the associated barcode-genotype pair. Due to the occurrence of template switching events during PCR amplification, one barcode might be assigned multiple genotypes to some extent. To improve the quality of barcode matching to specific genotypes, we applied a maximal parsimonious strategy to accept only one association supported by most PacBio reads or discarded the barcode if this strategy failed.

### NovaSeq sequencing and relative growth estimation

We performed paired-end 150 bp sequencing on each sample on the Illumina NovaSeq platform, with an estimated sequencing depth of 100, to obtain the frequency of each barcode within a sample. Barcode sequences of 30 nt were extracted from the sequencing reads, and the barcodes with nonidentical sequences from matching forward and reverse sequencing reads were excluded from further analysis. In addition, barcodes captured by PacBio technology were included in the downstream analysis. Barcode counts across technical repeats or biological replicates were combined. To ensure accurate estimation of the relative growth, 27,965 genotypes with a total of at least 100 reads from the three technical repeats at T0 were analyzed.

Relative growth at each culture stage of each competitive growth assay was calculated for individual mutations as [*f*_*t*_ (Mutant)/*f*_0_(Mutant)]/[*f*_*t*_ (WT)/*f*_0_(WT)], where *f*_*t*_ is the frequency of the barcode of a certain genotype in a post-competition sample; f_0_ is the frequency of the barcode of the same genotype at T0; *f*_*t*_(WT) is the frequency of the wild-type barcode in a postcompetition sample; and *f*_0_(WT) is the frequency of the wild-type barcode at T0 (Fig. S9).

To estimate the reliability of the relative growth measurements, we used genotypes with at least three barcodes to calculate the Signal-to-noise ratio (SNR). We calculated the observed standard deviation (*SD*) of the *f*_*t*_/*f*_0_ for different barcodes of each genotype. We then randomly perturbed the correspondence between genotype and barcode and recalculated the perturbed SD (*SD’*) of the *f*_*t*_ /*f*_0_ for different barcodes of each genotype. We repeated the perturbance 1,000 times to obtain 1,000 *SD’* values, whose average value was represented as 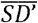. The SNR was then calculated as 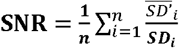, where n is the number of genotypes with at least three barcodes.

### Analysis of three-dimensional structure

The structure of MCR-1 in full length was modeled with Swiss Model. The EtpA [PDB: 5FGN] was used as a structural template, and the ribbon structure was presented with the PyMol software.

## Acknowledgment

This work was supported by the National Natural Science Foundation of China (grant number 81830103 and to G.-B. T. and J.-R. Y., grant numbers 82061128001, 81722030 to G.-B. T., grant numbers 31871320,32122022 to J.-R. Y., grant number 82002173 to S. F.), Guangdong Natural Science Foundation (grant number 2017A030306012 to G.-B. T.), National Key Research and Development Program (grant number 2017ZX10302301 to G.-B. T.), the National Key R&D Program of China (grant numbers 2021YFF1200904, 2021YFA1302500 to J.-R. Y.), Project of high-level health teams of Zhuhai at 2018 (The Innovation Team for Antimicrobial Resistance and Clinical Infection to G.-B. T.), and 111 Project (grant number B12003 to G.-B. T.);

## Author Contributions

Z. G., S. F., and L. L. contributed equally to this study. G.-B. T. and J.-R. Y. conceived the idea, designed and supervised the study. Z. G., S. F., L. L., J. L., L. Z. and H. Z. conducted the experiments, collected and analyzed the data. Z. G., S. F. and J.-R. Y. wrote the manuscript with inputs from all co-authors. All authors reviewed, revised, and approved the final report;

## Supplementary figures

**Fig. S1.**
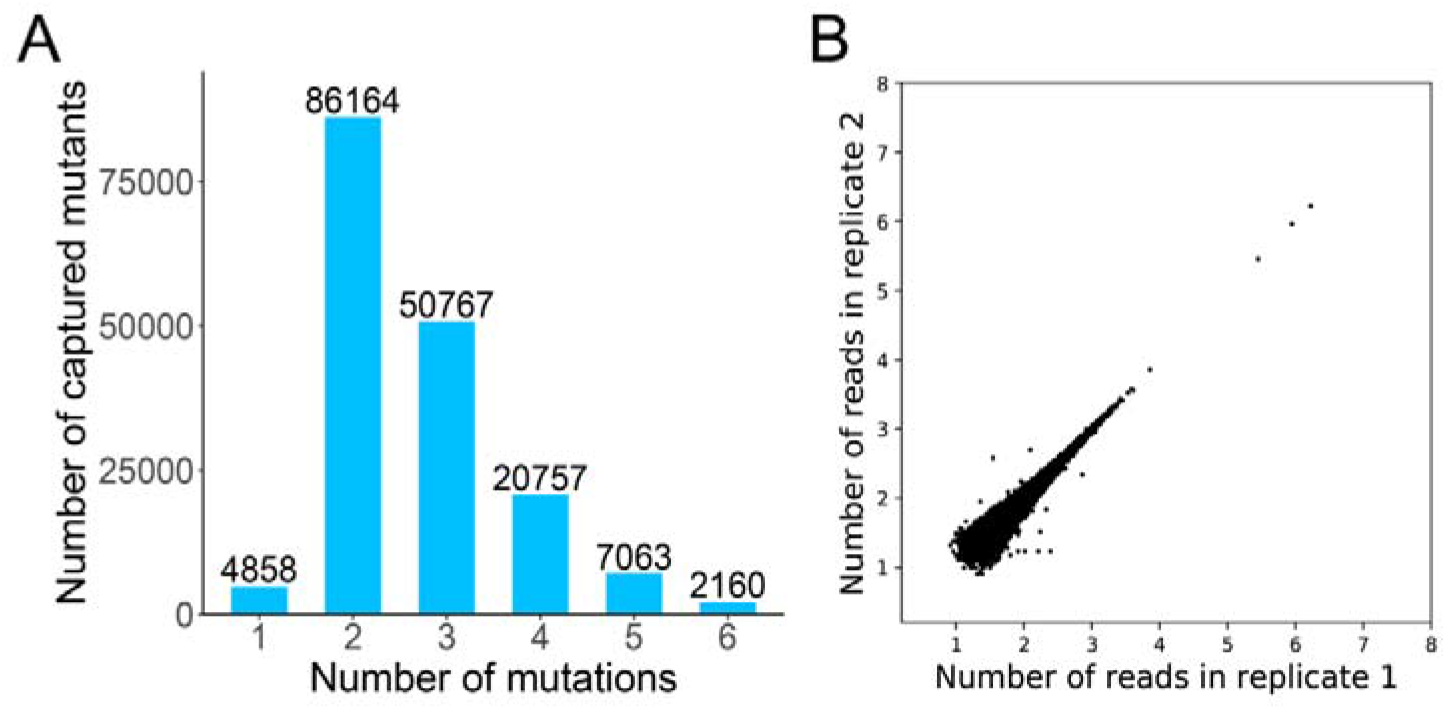
Quality of high throughput sequencing. **(A)** The number of genotypes captured by PacBio Sequel System. **(B)** Comparison of read pair numbers between two technical repeats at T0. Each dot represents a genotype, and only the genotypes with a total of ≥100 read pairs are considered. Pearson’s correlation coefficient (R) ≥ 0.99991.

## Reference

1 Liu, Y. Y. et al. Emergence of plasmid-mediated colistin resistance mechanism MCR-1 in animals and human beings in China: a microbiological and molecular biological study. Lancet Infect Dis 16, 161–168, doi:10.1016/S1473-3099(15)00424-7 (2016).

2 Shen, Z. Q., Wang, Y., Shen, Y. B., Shen, J. Z. & Wu, C. M. Early emergence of mcr-1 in Escherichia coli from food-producing animals. Lancet Infectious Diseases 16, 293–293, doi:Doi 10.1016/S1473-3099(16)00061-X (2016).

3 Feng, Y. J. Transferability of MCR-1/2 Polymyxin Resistance: Complex Dissemination and Genetic Mechanism. Acs Infect Dis 4, 291–300, doi:10.1021/acsinfecdis.7b00201 (2018).

4 Wang, R. B. et al. The global distribution and spread of the mobilized colistin resistance gene mcr-1. Nat Commun 9, doi:ARTN 117910.1038/s41467-018-03205-z (2018).

5 Shen, C. et al. Dynamics of mcr-1 prevalence and mcr-1-positive Escherichia coli after the cessation of colistin use as a feed additive for animals in China: a prospective cross-sectional and whole genome sequencing-based molecular epidemiological study. Lancet Microbe 1, E34–E43 (2020).

6 Wang, Y. et al. Changes in colistin resistance and mcr-1 abundance in Escherichia coli of animal and human origins following the ban of colistin-positive additives in China: an epidemiological comparative study. Lancet Infectious Diseases 20, 1161–1171, doi:10.1016/S1473-3099(20)30149-3 (2020).

7 Usui, M. et al. Decreased colistin resistance and mcr-1 prevalence in pig-derived Escherichia coli in Japan after banning colistin as a feed additive. J Glob Antimicrob Re 24, 383–386, doi:10.1016/j.jgar.2021.01.016 (2021).

8 Shen, C. et al. Prevalence of mcr-1 in Colonized Inpatients, China, 2011-2019. Emerg Infect Dis 27, 2502–2504, doi:10.3201/eid2709.203642 (2021).

9 Huang, H. et al. Colistin-resistance gene mcr in clinical carbapenem-resistant Enterobacteriaceae strains in China, 2014-2019. Emerg Microbes Infec 9, 237–245, doi:10.1080/22221751.2020.1717380 (2020).

10 Gao, R. S. et al. Dissemination and Mechanism for the MCR-1 Colistin Resistance. Plos Pathog 12, doi:ARTN e100595710.1371/journal.ppat.1005957 (2016).

11 Tietgen, M. et al. Impact of the colistin resistance gene mcr-1 on bacterial fitness. Int J Antimicrob Agents 51, 554–561, doi:10.1016/j.ijantimicag.2017.11.011 (2018).

12 Nang, S. C. et al. Fitness cost of mcr-1-mediated polymyxin resistance in Klebsiella pneumoniae. J Antimicrob Chemother 73, 1604–1610, doi:10.1093/jac/dky061 (2018).

13 Ma, K., Feng, Y. & Zong, Z. Fitness cost of a mcr-1-carrying IncHI2 plasmid. PLoS One 13, e0209706, doi:10.1371/journal.pone.0209706 (2018).

14 Yang, Q. et al. Balancing mcr-1 expression and bacterial survival is a delicate equilibrium between essential cellular defence mechanisms. Nat Commun 8, doi:ARTN 205410.1038/s41467-017-02149-0 (2017).

15 Feng, S. Y. et al. MCR-1-dependent lipid remodelling compromises the viability of Gram-negative bacteria. Emerg Microbes Infec 11, 1236–1249, doi:10.1080/22221751.2022.2065934 (2022).

16 Ramaloko, W. T. & Sekyere, J. O. Phylogenomics, epigenomics, virulome and mobilome of Gram-negative bacteria co-resistant to carbapenems and polymyxins: a One Health systematic review and meta-analyses. Environ Microbiol 24, 1518–1542, doi:10.1111/1462-2920.15930 (2022).

17 Smelikova, E., Tkadlec, J. & Krutova, M. How to: screening for mcr-mediated resistance to colistin. Clin Microbiol Infec 28, 43–50, doi:10.1016/j.cmi.2021.09.009 (2022).

18 Zhu, X. Q. et al. Impact of mcr-1 on the Development of High Level Colistin Resistance in Klebsiella pneumoniae and Escherichia coli. Front Microbiol 12, doi:ARTN 66678210.3389/fmicb.2021.666782 (2021).

19 Luo, Q. X. et al. In vitro reduction of colistin susceptibility and comparative genomics reveals multiple differences between MCR-positive and MCR-negative colistin-resistant Escherichia coli. Infect Drug Resist 12, 1665–1674, doi:10.2147/Idr.S210245 (2019).

20 Zhang, H., Zhao, D., Quan, J., Hua, X. & Yu, Y. mcr-1 facilitated selection of high-level colistin-resistant mutants in Escherichia coli. Clin Microbiol Infec 25, doi:10.1016/j.cmi.2018.12.014 (2019).

21 Gagliotti, C. et al. Reduction trend of mcr-1 circulation in Emilia-Romagna Region, Italy. Eur J Clin Microbiol 40, 2585–2592, doi:10.1007/s10096-021-04318-y (2021).

22 Chen, J. Z., Fowler, D. M. & Tokuriki, N. Environmental selection and epistasis in an empirical phenotype-environment-fitness landscape. Nat Ecol Evol 6, 427–438, doi:10.1038/s41559-022-01675-5 (2022).

23 de Visser, J. A. G. M., Elena, S. F., Fragata, I. & Matuszewski, S. The utility of fitness landscapes and big data for predicting evolution. Heredity 121, 401–405, doi:10.1038/s41437-018-0128-4 (2018).

24 Feng, S. Y. et al. Prediction of Antibiotic Resistance Evolution by Growth Measurement of All Proximal Mutants of Beta-Lactamase. Mol Biol Evol 39, doi:ARTN msac08610.1093/molbev/msac086 (2022).

25 Gao, R. et al. Dissemination and Mechanism for the MCR-1 Colistin Resistance. Plos Pathog 12, e1005957, doi:10.1371/journal.ppat.1005957 (2016).

26 Puchta, O. et al. Network of epistatic interactions within a yeast snoRNA. Science 352, 840–844, doi:10.1126/science.aaf0965 (2016).

27 Green, M. R. & Sambrook, J. The Hanahan Method for Preparation and Transformation of Competent Escherichia coli: High-Efficiency Transformation. Cold Spring Harb Protoc 2018, doi:10.1101/pdb.prot101188 (2018).

28 Chaisson, M. J. & Tesler, G. Mapping single molecule sequencing reads using basic local alignment with successive refinement (BLASR): application and theory. Bmc Bioinformatics 13, doi:Artn 23810.1186/1471-2105-13-238 (2012).

29 Wenger, A. M. et al. Accurate circular consensus long-read sequencing improves variant detection and assembly of a human genome. Nat Biotechnol 37, 1155-+, doi:10.1038/s41587-019-0217-9 (2019).

