## Supplementary material for "The fitness landscape of the mobilized colistin resistance gene *mcr-1*": Table S4

**Table S4 MIC value of antibiotics for BW25113 carrying wild-type MCR-1 or variant**

| Antibiotic | MIC (μg/ml) | | | | | | |
| --- | --- | --- | --- | --- | --- | --- | --- |
|  | MCR-1 | C584T | A557G | A110C | T5C | G577T | T18G |
| Colistin | 16 | 16 | 16 | 16 | 16 | 16 | 16 |
